# Megaherbivores modify forest structure and increase carbon stocks through multiple pathways

**DOI:** 10.1101/2021.12.23.473993

**Authors:** Fabio Berzaghi, François Bretagnolle, Clémentine Durand-Bessart, Stephen Blake

## Abstract

Megaherbivores have pervasive ecological effects. In African rainforests, elephants can increase aboveground carbon, though the mechanisms are unclear. Here we combine a large unpublished dataset of forest elephant feeding with published browsing preferences totaling > 120,000 records covering 700 plant species, including nutritional data for 102 species. Elephants increase carbon stocks by: 1) promoting high wood density tree species via preferential browsing on leaves from low wood density species, which are more digestible; 2) dispersing seeds of trees that are relatively large and have the highest average wood density among tree guilds based on dispersal mode. Loss of forest elephants could cause a 5-12% decline in carbon stocks due to regeneration failure of elephant-dispersed trees and an increase in abundance of low wood density trees. These results show the major importance of megaherbivores in maintaining diverse, high-carbon tropical forests. Successful elephant conservation will contribute to climate mitigation at a scale of global relevance.

## Introduction

Megaherbivores (body mass > 1000 kg) can have profound effects on vegetation, carbon stocks, and nutrient cycling^1–3^. However, knowledge on the ecosystem role of megaherbivores comes predominantly from African savanna ecosystems^1,4^. In tropical forests, initial evidence suggests that these large herbivores might also have profound effects^5–7^. Until the Late Pleistocene, tropical forests hosted a wide variety of megaherbivores^1^. Today, the Asian (*Elephas maximus*) and African forest elephant (*Loxodonta cyclotis*) are the only forest-dwelling megaherbivores with extensive ranges and functionally unique characteristics: large size, diverse behaviors, and highly varied diets. Examples of “ecosystem engineering” have been observed in forest elephants (“elephants”) through seed dispersal^6,8^ and disturbance^9^ (i.e., browsing and trampling). By reducing tree density, elephants promote the growth of larger trees with consequent drop in light and water availability in the understory. As a result, forests with elephants hold more aboveground carbon (AGC) because of a greater abundance of large late-successional tree species which have high wood density (WD)^5^. Berzaghi et al. (2019) evaluated the effect of elephants in terms of a generic disturbance-induced mortality from trampling and uprooting. However, megaherbivores interactions with ecosystems happen also via more delicate processes such as herbivory and seed dispersal^6,10,11^, whose influence on forest structure and AGC is currently unknown. The high daily food consumption (100-200 kg^12^) and broad diet (over 350 species^13^) of elephants suggest that feeding preferences could drive shifts in tree species composition by promoting growth and survival of less-desirable browse species. Folivores prefer leaves high in protein and minerals and low in fiber and chemical defenses (e.g., tannins) ^14^. Among woody plants, the abundance of defensive chemicals and non-digestible fiber are positively correlated to WD because slow-growing species invest more in structural and chemical defenses than faster growing species^15,16^. We hypothesize that elephants promote high AGC by preferentially browsing leaves from low WD trees.

We also investigate the connection between elephant-dispersed trees (“Obligate” trees *sensu*^6^) and AGC. Large-seeded animal-dispersed trees have relatively large diameters, high WD, and contribute significantly to AGC^17^. Forest elephants are prodigious seed dispersers, moving more seeds from more species than any other animal^6^, but the contribution of Obligate trees to forest structure and AGC has not been evaluated. We hypothesize that the combined effects of elephant browsing, which *decreases* fitness of preferred food species, and seed dispersal, which *increases* fitness of dispersed species, are likely to have profound effects on forest structure and AGC. If supported, these two hypotheses would confirm the ecosystem role of elephants in promoting high carbon stock forests by increasing the fitness of large, high WD trees^5^. To test these hypotheses, we combined forest inventories and feeding data collected in Ndoki (Republic of Congo) and LuiKotale (Democratic Republic of Congo) with published elephant diet selection data across the Afrotropics. We analyzed WD as a function of elephant browsing preferences and the nutritional properties of leaves to investigate the mechanisms driving elephant choices and the influence of these choices on AGC. We then synthesized, based on literature, quantitative measures of the effects of elephants on forest properties and processes and schematically organized these findings. This synthesis supports our hypotheses, identifies research gaps, and provides input for modeling elephants using statistical and process-based models. Our results greatly enhance our understanding of the contribution of elephants to forest functioning and are key to evaluate the consequences of past megaherbivore extinctions and to inform conservation and management policy.

## Results

### Forest elephants browse most frequently on low wood density species

We collated forest elephant feeding data from eight different sites across tropical Africa: West (N = 4), Central (2), and East (2). The global dataset (collection of all sites) includes 197,557 sampled plants from 702 plant species for which WD could be determined (Data Table S1). The actual number of sampled plants is higher because three studies did not report their total sample size (Table S1). All sites, except Bia National Park (NP) and Santchou Wildlife Reserve, recorded feeding preferences by accounting for the abundance of elephant-selected species. In those two sites, limited information was available on the relative abundance of elephant-preferred species^18,19^. Feeding preference metrics were similar across sites and could be assimilated into three groups indicating high, intermediate, and low preference (Methods). Globally, feeding preference was negatively correlated with WD (Fig 1). This trend was confirmed even when excluding Ndoki, the only site where both the number of feeding events and their quantity were recorded (Fig. S1). In seven out of eight sites the low preference group had the highest WD compared to the intermediate and high preference groups (Fig. 2). In half of the sites the WD average of the low preference group was significantly lower than the other groups (Fig. 2). Only in Bia NP this trend was not observed. Here, the medium preference group had the highest WD and the low preference group had the lowest (see discussion for possible explanation). The WD of trees dispersed by elephants was higher compared to highly browsed species in four out of five sites, but only two were significantly different (Fig. 2). This is compatible with the hypothesis that elephants facilitate species with higher WD through dispersal and browsing.

**Fig. 1.**
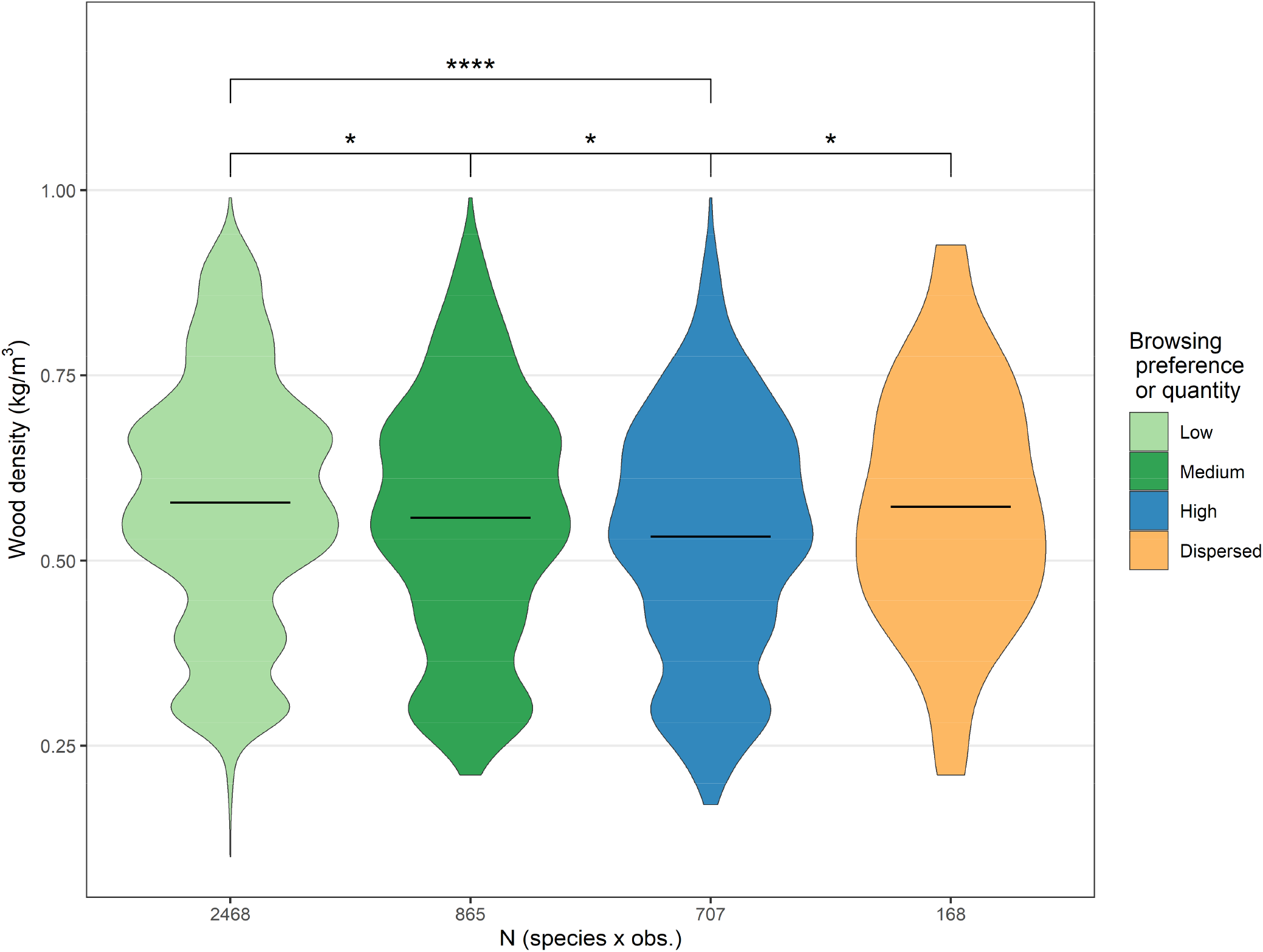
Wood density of elephant-dispersed and browsed trees by preference across tropical Africa. Elephants prefer to browse on low wood density trees and disperse seeds from trees with high wood density. The x-axis indicates the number of species plus observations (only at Ndoki). The elephant-dispersed group includes all tree species of which seeds were dispersed by elephants and includes elephant-obligate and non-obligate (dispersed by elephants and other animals). Significance level of pairwise statistical comparison: ^*^P < 0.05; ^****^P < 0.0001.

**Fig. 2.**
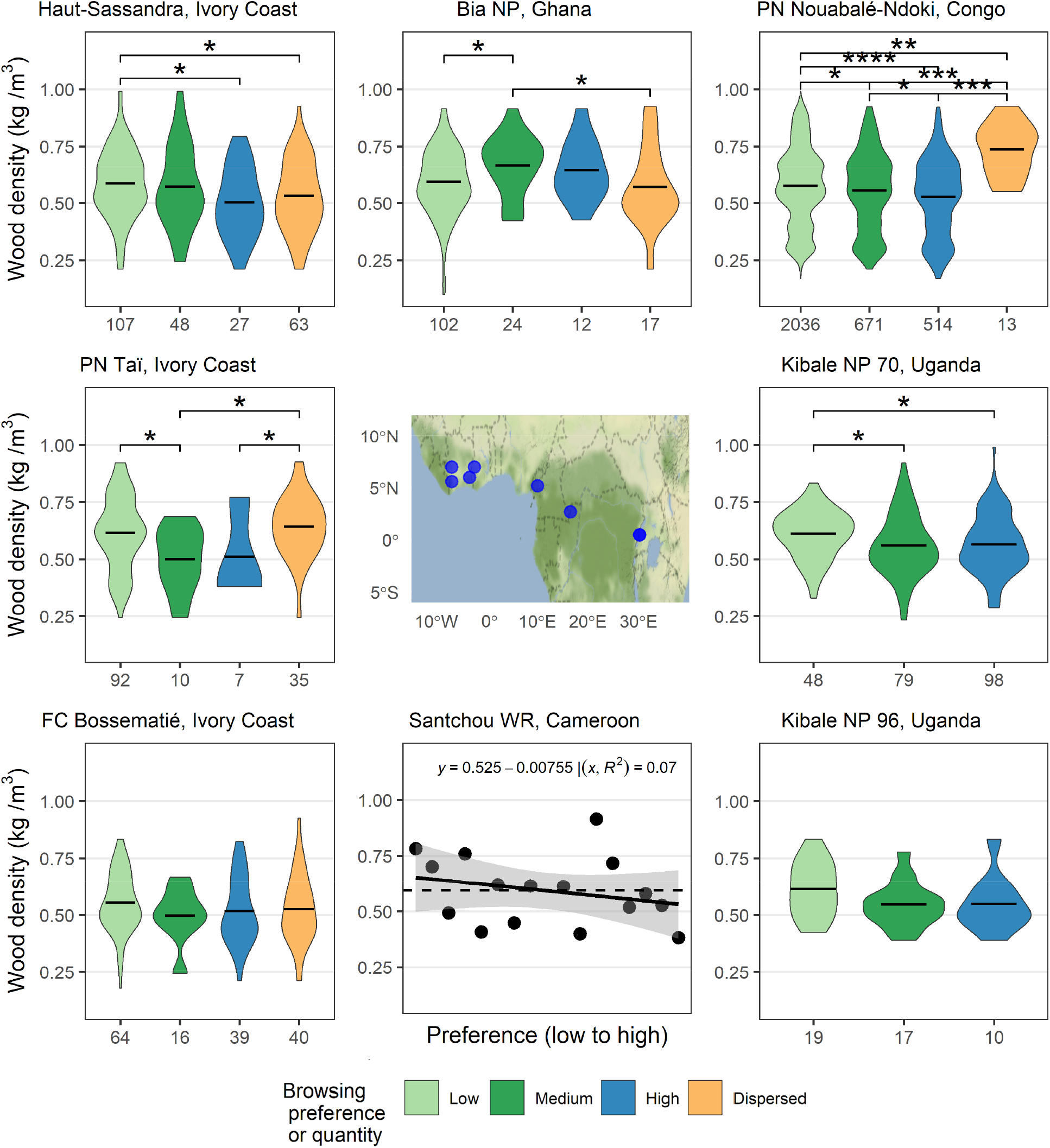
Wood density of elephant-dispersed trees and across elephant browsing preferences groups in eight sites in tropical Africa. The x-axis indicates the number of species in each group, or the number of observations at Ndoki. In Santchou only ordinal ranking was provided. The elephant-dispersed group includes all tree species of which seeds were dispersed by elephants and includes elephant-obligate and non-obligate (dispersed by elephants and other animals). Significance level of pairwise statistical comparison: ^*^P < 0.05; ^**^P < 0.01; ^***^P < 0.001; ^****^P < 0.0001.

Our data from Ndoki (understory and overstory) and LuiKotale (overstory) showed that correlation between WD and plant abundance is not strong; at most, high WD species are slightly more abundant than low WD ones (Fig. S2-S3). Elephants made specific choices regardless of the abundance of species. For example, in Ndoki understory vegetation plots, *Rinora welwitschii* and *Diospyros bipindensis* were recorded 559 and 468 times respectively (from a dataset of 6548 tree stems from at least 151 species, Methods). Yet, of 5458 feeding events, only two involved *D. bipindensis* and *R. welwitchii* was never browsed. A comparison of mean WD of the top 10 most frequently browsed species that were not present in plots with that of the top 10 most frequent species in plots which were never browsed revealed that browsed species have significantly lower WD than non-browsed species (ANOVA, P=0.05).

### Nutritional properties of elephant food

We investigated the mechanisms driving browsing preferences using a global database of plant nutritional values^20^. We analyzed WD as a function of leaf and fruit nutritional properties. The data covered 102 plant species and 807 records of essential biomolecules (crude protein, minerals, carbohydrates, fiber, and nonstructural carbohydrates), structural and defensive compounds (tannins, lignin, and fibers), which reduce food palatability and digestibility (% of assimilated food). Wood density appears to decrease as leaf protein, minerals, and digestibility content increase, while fibers and tannins are highest at intermediate values of WD (Fig. S4); however, the only significant correlations were for fibers (R^2^ = 0.14, P < 0.001), dry matter digestibility (R^2^ = 0.1, P = 0.036), and tannins (R^2^ = 0.16, P = 0.011) (Fig. S1). Analysis of leaf properties across browsing preference groups showed dry matter digestibility (R^2^ = 0.11-0.20, P > 0.05) and fibers (R^2^ = 0.16-0.26, P <0.001) were highest at intermediate WD levels (Fig. 3a); no other appreciable correlations were detected among the other nutritional properties (Fig. S5). Highly preferred species had a narrower and lower range of fibers and tannin which is reflected into a higher digestibility compared to the other groups (Fig. S6). Proteins, carbohydrates, and hemicellulose were also higher in the high-preference group, but no statistical differences were detected in part because of the low sample size for saccharides (Fig. S6). High WD seems to be correlated with high values of leaf carbohydrates and fat with the exception of lignin and non-structurally carbohydrates (Fig. S4); however, the sample size for these properties was small (n = 7). This suggests that leaves from low WD plants might be less nutritious but more digestible because of their lower fiber and tannin content compared to high WD plants. Elephants, like most large herbivores, must balance digestibility and food quality in their food choices, but more data would be needed to draw more conclusive results on this trade-off.

**Fig. 3.**
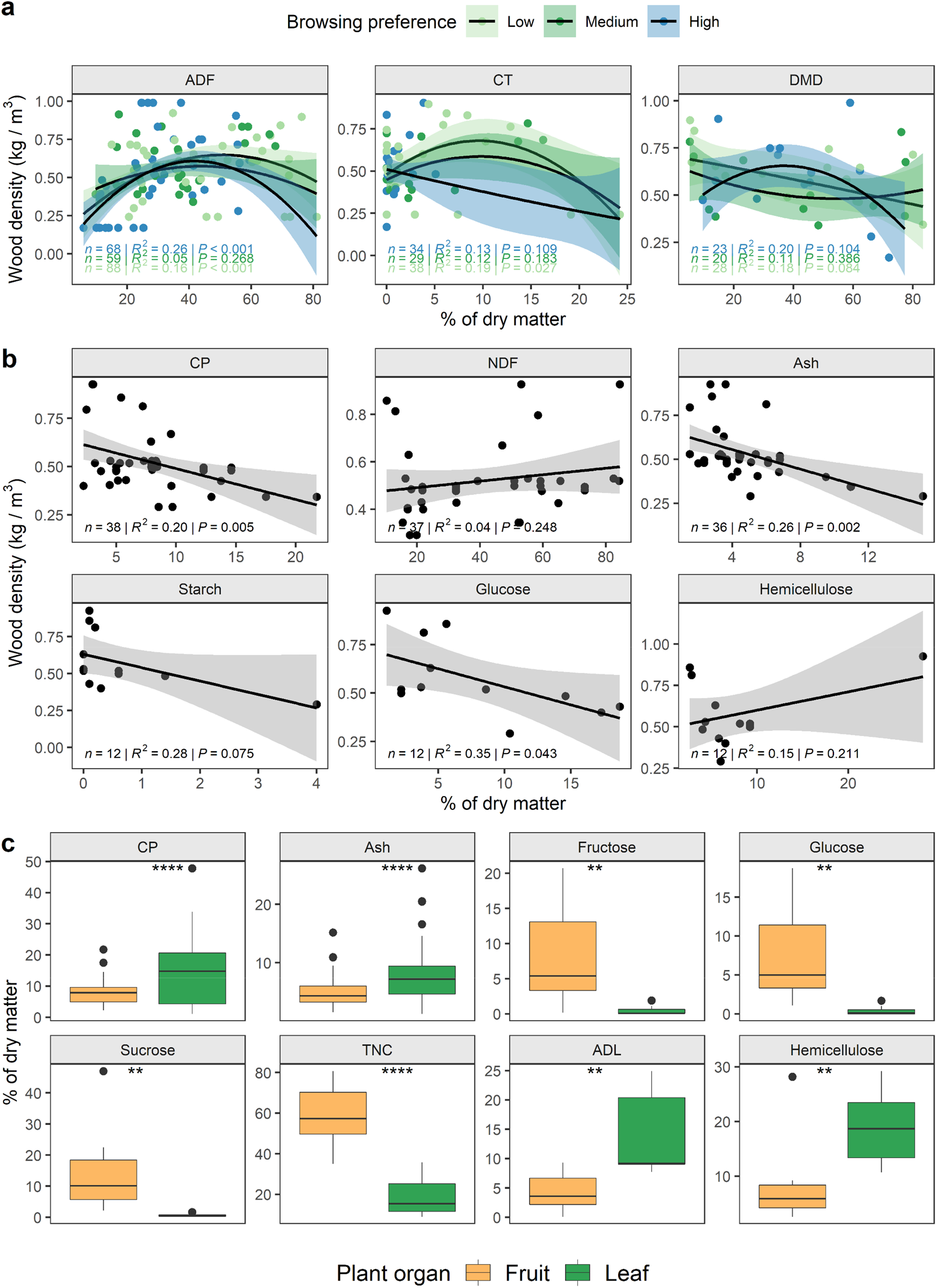
Comparison of leaf and fruit nutritional properties and relations with wood density. ADF = acid detergent fiber, ADL = acid detergent lignin, Ash = minerals, CT = condensed tannins, CP = crude protein, DMD = dry matter digestibility, NDF = neutral detergent fiber, TNC = total non-structural carbohydrates. (a) wood density as a function of leaf properties of plants based on browsing preference; (b) wood density as a function of fruit dispersed by elephants. These include Obligate and Non-Obligate trees (see text); (c) comparison of nutritional properties between fruit and leaves. Significance level of pairwise statistical comparison: ^**^P < 0.01; ^****^P < 0.0001.

Fruits produced by high WD trees are high in protein, minerals, starch, and glucose and low in sucrose and cellulose (Fig. 3b). We found no significant correlation between WD and the other properties (Fig. S7). This implies that fruit from high WD trees are highly nutritious and likely therefore to be preferred by elephants and other frugivores compared to fruit from low WD trees. When comparing fruits and leaves, we found, as expected, that leaves are higher in protein (P <= 0.0001), minerals (P <= 0.0001), lignin (P <= 0.01), and cellulose (P <= 0.01), while fruits have higher nonstructural carbohydrates (P <= 0.0001) and sugars (P <= 0.01) (Fig. 3c); leaf and fruit other properties were not statistically different (Fig. S8). Thus, fruit provide short-term usable energy, but leaves contain nutrients for longer-term physiological processes. This might act as a physiological limitation and an evolutionary bottleneck of obligate frugivores to develop very heavy body mass. Overall, these correlations suggest that elephant preference for leaves from lower WD species is due to their higher digestibility (i.e., lower fiber and tannin content), while fruits from high WD species are preferred because of their high content of protein, starch, glucose, and minerals and low cellulose.

### Elephant-dispersed trees are larger and have higher wood density compared to trees with other dispersal modes

We identified five dispersal modes in Ndoki and LuiKotale: Gravity/dehiscence (GD), wind, elephants and other animals (Non-Obligate), elephants (Obligate), and Other-Animals^6,21^ (total of 238 species, complete list in Supplementary Data). The analysis of the variation of WD as a function of dispersal mode revealed that Obligate species had the highest average WD in both sites (Fig. 4a). In LuiKotale, only GD and Obligate species are statistically different than Non-Obligate (P ≤ 0.05). Non-Obligate (both sites) and wind-dispersed (Ndoki) species have the lowest WD. Obligate (both sites), GD (Ndoki), and Wind (LuiKotale) are the dispersal modes with the lowest number of species.

**Fig. 4.**
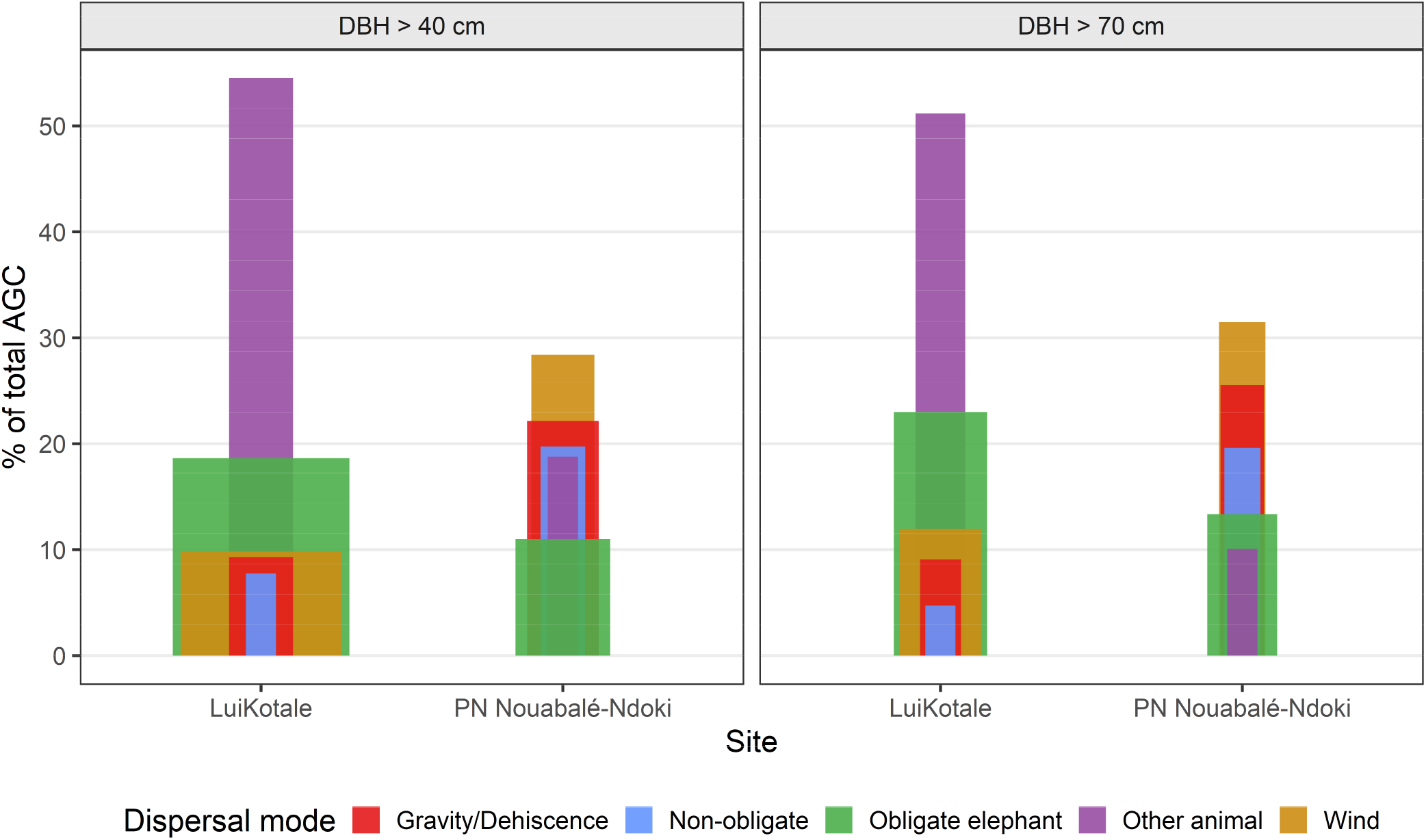
Relative contribution of dispersal guilds to aboveground carbon at different size thresholds. The bar width is an indication of the relative importance of each guild for AGC in relation to the total number of stems in the forest. It is calculated for each guild by dividing the percentage of total AGC for the percentage of stems at each site. Larger ratios (wider bars) indicate a large contribution to AGC by a small number of stems.

The distribution of stem size classes across dispersal mode was mostly similar at the two sites (Fig.4b). Obligate and wind-dispersed tree communities are characterized by few smaller trees, a higher number of larger trees, and are overrepresented in the 125-250 cm range compared to trees with other dispersal modes (Fig. 4b). Obligate trees represent the largest proportion in LuiKotale, 35% and 55%, and second largest in Ndoki, 26% and 28%, of stems with diameter > 150 and 175 cm, respectively. They also comprise the second (LuiKotale, 30%) and third largest (Ndoki 16%) proportion of stems with diameter > 125 cm. The proportion of wind-dispersed trees also increases with size class. Non-Obligate and Other-Animals trees are most abundant in the lower size classes between 40-125 cm (Fig. 4b). This distribution of stem size across dispersal modes might reveal the long-term history of these forests emerging after the recolonization of savannas.

**Fig 3.**
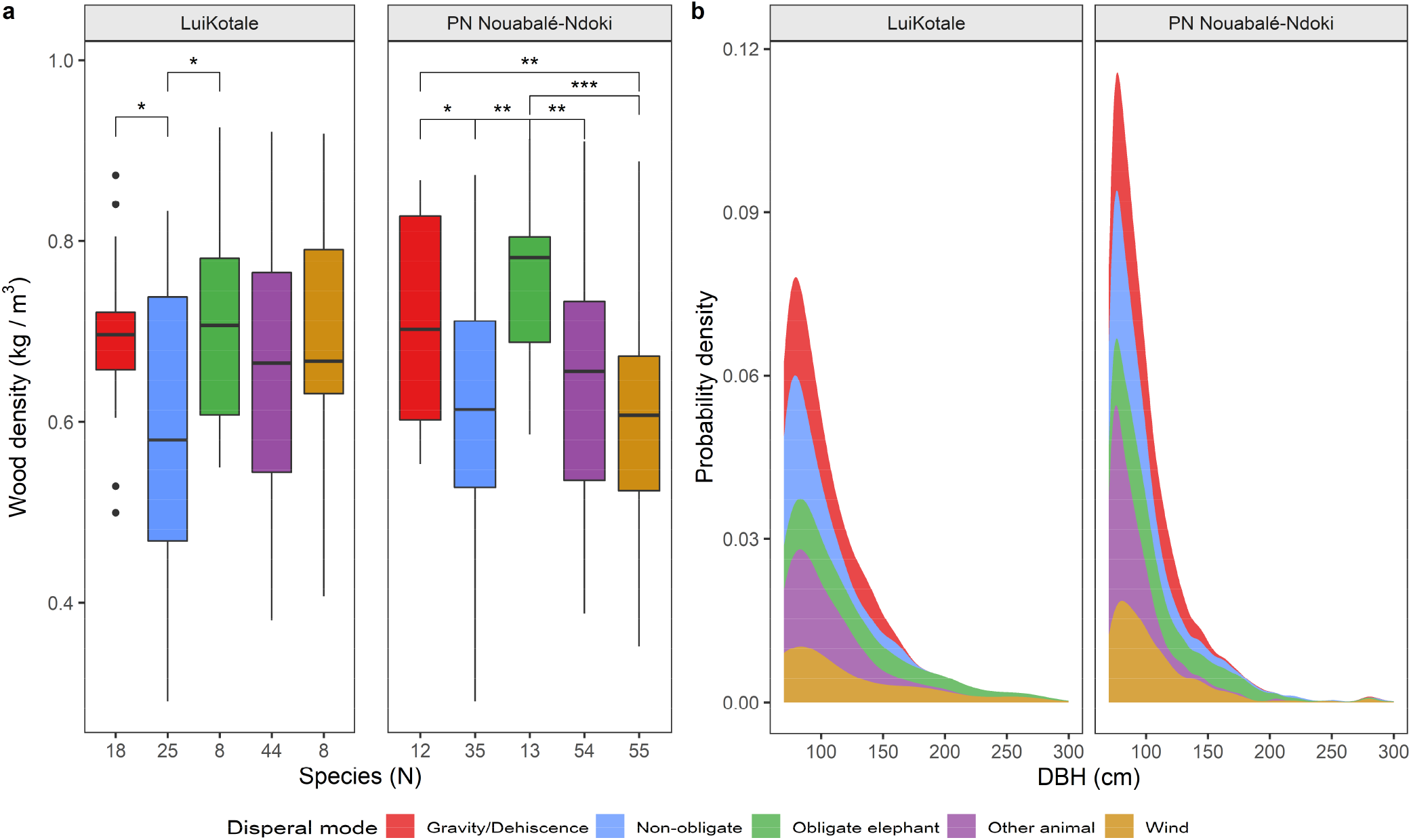
Properties of tree species and forest structure at Ndoki and LuiKotale. (a) Variation in wood density as a function of dispersal mode. Significance level of pairwise statistical comparison: ^*^P < 0.05; ^**^P < 0.01; ^***^P < 0.001. (b) Kernel density distribution of trees according to their dispersal modes at Ndoki.

### Contribution of elephant-dispersed trees to aboveground carbon

The distribution of aboveground carbon in trees (diameter ≥ 40 cm) grouped by dispersal mode reveals diverse patterns in Ndoki and LuiKotale (Fig. 5). In Ndoki, AGC is more evenly distributed among dispersal modes. Abiotically-dispersed trees account for ∼50% and Obligate for ∼11% of AGC (smallest biomass pool). In LuiKotale, trees dispersed by other animals store 54% of AGC and ∼19% is stored in Obligate trees (second largest biomass pool). At Ndoki, our sampling of vegetation was biased toward the monodominant *Gilbertiodendron dewevrei* forest, which occupies a proportion of Ndoki^13^ by following watercourses, as do forest elephants. If *G. dewevrei* forest was removed from the analysis, and only mixed species *terra firma* forest considered, the contribution of Obligate species would be profound. When considering only larger trees (diameter ≥ 70 cm), a higher percentage of AGC is stored in Obligate (23% LuiKotale and 13% Ndoki) and abiotically-dispersed (57% Ndoki) trees (Fig. 5). Notably, at both sites the few Obligate species have the highest relative contribution to AGC despite their low stem count (i.e. highest AGC to stem ratio represented by bar widths in Fig. 5). This is explained by their high WD and high relative abundance in the large size classes (diameter > 125 cm) (Fig. 3b). The loss of forest elephants might greatly diminish or prevent the recruitment of future Obligate trees in addition to affecting Non-Obligate species. If we substitute Obligate trees with random trees with other dispersal modes proportionally to their relative abundance (Methods), the loss of AGC was estimated to be 12% (s.d. ± 1.2) at LuiKotale and 5% (s.d. ± 0.4) at Ndoki. Thus, the “other” trees cannot completely compensate the contribution of Obligate trees. The important role of large trees in AGC^22,23^ and the widespread decline of forest elephants make the plight of Obligate species critical for the future of AGC in African tropical forests.

### Ecological processes influenced by elephants

The effects of savanna elephants on their environment have been heavily studied^24^. Forest elephants seem to perform similar actions to their savanna counterparts but few studies have quantified their impacts. We synthesized the literature and selected studies that provided quantifiable measures of the mechanisms of ecosystem engineering by elephants expressed in terms of rates, equations, or data. Of all the possible ecological processes influenced by elephants^25^, only a few have been quantified and most of them only once or twice. Many other studies exist on seed dispersal or browsing preferences but we could not quantify or generalize with equations their consequences on ecosystem properties. Savanna and forest elephants alike topple small trees to access foliage, scar and debark trunks but the impacts of these foraging effects on mortality in forests are poorly quantified (Table 1). Data on debarking and scarring and forest properties (forest openness, stem density, AGC, and WD) come from single studies (Table 1). Only one study quantified forest properties as a function of elephant trails^26^. However, a few general conclusions can be drawn from our synthesis. The mortality rate inflicted by elephants to large trees (DBH >10 cm) is between 1-2% which is similar to the background mortality of African tropical forests^27^. The mortality of seedling and saplings is several times higher compared to large trees. Distance from trails is a key parameter when assessing the effect of elephants on forest properties. There is also a clear relationship between canopy openness, reduced regeneration, and elephant preference, however this has not been estimated in more quantitative terms such as visitation frequency or biomass consumption. Less robust conclusions can be drawn on forest elephant impacts on the density of small trees and the mortality rate of large trees due to debarking. Elephant density (individuals/km^2^) should be accounted for when evaluating the magnitude of elephant effect on forest properties and processes, particularly for comparisons across studies or sites. Densities were not always reported and we highlight the need for considering this parameter when extrapolating results to other areas. We also suggest that studies should report the equations of fitted regressions, which would be useful for modelling approaches.

**Table 1.** Summary of literature review of the ecological effects of elephants in closed canopy forests across the Afrotropics. Only studies that provided a quantitative measure or a mathematical function were included in the table. DBH = diameter at breast height

| Description | Quantitative result | Qualitative result (if any) | Location, elephant density, and sampled area | Ref. |
| --- | --- | --- | --- | --- |
| *<u>Mortality - regeneration</u>* |  |  |  |  |
| Mortality rate after elephant damage (DBH > 10 cm) | 1.4% (Annual rate) |  | Kibale NP, Uganda, 5.3 ha | 28 |
| Recovery rate after elephant damage (DBH > 10 cm) | 1.2% (Annual rate) |  | Kibale NP, Uganda, 5.3 ha | 28 |
| Sapling mortality rate | 4% (Annual rate) |  | Kibale NP, Uganda, logged | 29 |
| Seedling and saplings mortality (height > 10 cm) | 15-18% |  | Kibale NP, Uganda, logged | 30 |
| Tree toppling & branch breaking | 2 - 9.9 cm DBH:<br>- toppled 40.9%<br>- broken branch 24%<br>> 10 cm DBH<br>- toppled 6.9%<br>- broken branch 7% | Tree toppling and broken branches decline sharply for trees > 10 cm DBH. Larger trees suffer more bark stripping | Bwindi NP, Uganda, 0.97 ha | 31 |
|  | 68% breaks by elephants | Most breaks between 1 m and 3 m height, 2 cm and 6 cm DBH | Several sites, Gabon | 9 |
| Reduced regeneration | - Browsed species contained 19% saplings of canopy and 48% subcanopy species<br>- Trampling, movement, and grubbing prevents regeneration in 25% of the sampled area |  | Shimba Hills National Reserve, Kenya (both forest and savanna elephants common in the part) | 32 |
|  | - Canopy opening < 20% and forest gaps < 300 m<sup>2</sup> reduces elephant |  | Kibale NP, Uganda | 33 |
| *<u>Forest properties</u>* |  |  |  |  |
| Mean DBH from trail (distance from trail) | 52 cm (0-5m)<br>23 cm (21-25m) | Mean DBH decreases away from trails | Salonga, 0.05 ind/km<sup>2</sup>, 100 km of transects | 26 |
| Understory openness & elephant encounter rate | $y = 0.2386x + 0.055$ | Dung encounter rate increases linearly with understory openness | Salonga, 0.05 ind/km<sup>2</sup>, 100 km of transects | 26 |
| Tree species composition & distance from trail |  | Distribution of fruit-preferred and browse-preferred trees varies as a function of distance from trails | Salonga, 0.05 ind/km<sup>2</sup>, 100 km of transects | 26 |
| Seedling and sapling density and damage near elephant trees | Elephant presence increases chances of damage to seedlings (84%) and saplings (24%) |  | Ivindo NP, Gabon | 34 |
| Aboveground carbon | $y = -0.0841 + 0.3311x - 0.0630x^2$ | Percentage change in aboveground carbon (y) as a function of elephant density (x) | Process-based vegetation model | 5 |
| Stem density | Reduced density of plants between < 1 cm and ≥ 1 m in height |  |  | 9 |
| Stem scarring (DBH > 10 cm) | 16% of stems scarred |  | Rabongo, Uganda, 7ha | 35 |
| Debarking height | Species-specific results | Percentage of debarked trees, average diameter and debarking height |  | 18,19 |

## Discussion

We have shown that, across their range, forest elephants browse most frequently on woody species with low WD (Fig 1-2). Some abundant species were completely ignored while many rare species were frequently consumed, though elephants had local preferences at the species level. The exception of Bia NP might be due to woody lianas and climbers being particularly common both in the forest and in elephant diets compared to the other sites^18^. Elephant browsing preferences are likely driven, in part, by leaf nutritional properties. Low WD and frequently-browsed trees produced more digestible leaves contain less fibers and tannins. This strongly supports our hypothesis that elephant browsing increases the AGC of central African forests by reducing the fitness of preferred fast-growing species, which promotes survival and growth of slow growing, high WD species. A previous study also suggested that if elephants are extirpated from forests, the community average WD declines as fast growing species become more abundant^5^. Without elephants, the forest average WD will become lower but by how much is difficult to quantify.

Elephants also influence AGC by dispersing seeds of tree species having high WD which are also over-represented in large sizes (Fig. 4). The reason for a higher relative abundance of Obligate trees in larger size classes is unclear, but may be due to the combination of life history traits of large seeded species and forest succession history. Wood density is correlated with structural strength, low mortality, and resistance to decay which favor large size and longevity (though slow growth means that attaining large size takes longer for these species)^36,37^. However, the strength of this correlation declines with increasing tree size^38^ and some of the largest trees in the forest are fast growing, wind dispersed species of low WD (e.g. *Triplochiton scleroxylon* and *Ceiba pentandra*). Whatever the underlying reasons for their large size and high WD, Obligate trees contribute significantly to AGC (Fig. 5). Declines in abundance or the complete extirpation of forest elephants will therefore reduce recruitment through extinction of elephants^6^ and result in an important reduction in AGC, estimated at 5-12% at our two study sites. Many other Non-Obligate tree species might also experience reduced recruitment rates because elephants contribute to a large proportion of their seed dispersal^6^. The scale of such declines in AGC would depend on the specifics of tree community dynamics, which can also be predicted with long-term model simulations^39^.

The current knowledge base on the processes and properties of forest that are influenced by elephants is better developed in some areas, such as small plant mortality (Table 1). However, many processes and properties have received less attention or have been evaluated in terms that cannot be fully employed to model the effects of elephants on forests. There is a lack of repeated experiments in different sites to verify if locally observed effects are consistent across sites and to evaluate relation between elephant density and the magnitude of the change. Yet, the current knowledge provides a good starting point to better characterize elephant effects in modeling studies.

Our results add further evidence that large and mega-herbivores contribute to enhance AGC in tropical forests through different mechanisms. Until the late Pleistocene, many large herbivores inhabited Amazonian and southeast Asian tropical forests and could have had a significant effect on the functioning of those ecosystems. Today, the disappearance of forest elephants will result in loss of AGC between 5-12% due to the decline of Obligate trees. An additional decline in AGC is also expected as the forest shifts to a higher abundance of low WD plants. Process-based vegetation models based on these findings and the processes shown in Table 1 will help better estimate the long-term consequences of elephants decline or, hopefully, repopulation^39^. The significant contribution of forest elephants to carbon stocks and biodiversity should be accounted to prioritize conservation of the species and their habitat, and implemented in climate change mitigation policy, and leveraged to promote and finance nature-based solutions in tropical Africa.

## METHODS

### Study sites

The Ndoki Forest (“Ndoki” 1.5–3° N, 16–17° E) lies in the northern Republic of Congo. The climate is transitional between the Congolo-Equatorial and sub-equatorial zones with a mean annual rainfall of ca. 1400 mm (Ndoki Forest records) ^6,13^. Topography varies from *terra firma* uplands and flat plateaus to the northwest to the extensive floodplain of the Likouala aux Herbes River to the southeast. Soils are of three main types: arenosols to the north and west, ferrasols to the southeast in the Likouala aux Herbes basin on *terra firma*, and gleysols in the swamps further southeast. Ndoki is embedded in wet Guineo-Congolian lowland rainforest, transitioning to the north into dry Guineo-Congolian lowland rainforest, and into swamp forests to the south. *Terra firma* is dominated by Sterculiaceae-Ulmaceae semi-deciduous forest, with large patches of mono-dominant *Gilbertiodendron dewevrei* forest along watercourses and upland plateaus, and swamp forests^13^. The Ndoki fauna includes several large charismatic species such as forest elephants, western lowland gorillas (*Gorilla gorilla gorilla*), common chimpanzees *(Pan troglodytes troglodytes*), forest buffalo (*Syncerus caffer nanus*), bongo (*Tragelaphus eurycerus*), and leopards (*Panthera pardus*). The human population density is low (<1 inhabitant/ km^2^) and the immediate study area contains no permanent human settlement.

The LuiKotale research site (LK) is located within the equatorial rainforest (2°470S, 20°210E), at the south-western fringe of Salonga National Park in the Democratic Republic of the Congo^21^. The study site covers >60 km2 of primary evergreen lowland tropical forest. The climate is equatorial with abundant rainfall (>2000 mm/yr) and two dry seasons, a short one around February and a longer one between May and August. Mean temperature at LuiKotale ranges between 21 °C and 28 °C with a minimum of 17 °C and a maximum of 38 °C (2007–2010). Two major habitat types can be distinguished. The dry (terra firma) forest and the wet temporarily and permanently inundated forest. The dry habitat dominates with heterogeneous species composition covering 73% and patches of mono-dominant *Gilbertiodendron* spp. covering 6% of the site. The wet habitat consists of heterogeneous forest temporarily (17%) and permanently (4%) inundated^21^. The LuiKotale fauna includes several large species such as elephants (almost extinct), bonobos (*Pan paniscus*), forest buffalo, bongo (*Tragelaphus eurycerus*), and leopards (*Panthera pardus*). Similarly to Ndoki, the human population density is low (<1 inhabitant/ km^2^) and the immediate study area contains no permanent human settlement.

### Elephant food selection

Fresh elephant trails were followed opportunistically over the course of two years in Ndoki across a range of habitat types including permanent swamps, seasonally inundated forests, and *terra firma* open and closed canopy forest. In the case of woody species, a single feeding event was defined as all fresh feeding signs on an individual plant, regardless of plant parts consumed, though all parts consumed were also recorded. At each feeding site data were collected on location (using a handheld Garmin GPS) estimated age (fresh [<24 hrs] or recent [24-48hrs]), plant species, plant part consumed (leaf, stem, bark, wood, roots, etc.), estimated amount consumed on a 1-4 scale (rare, few, moderate, and abundant). The total feeding events recorded were 4941.

Over a 3-yr period throughout the Ndoki Forest, the seed content of 855 piles of fresh intact elephant dung was quantified. Dung piles were broken apart with sticks, and fibers were thoroughly teased apart. In each dung pile, all seeds were identified to species, and all seeds >1 cm on the longest axis were counted.

### Analysis of elephant diet

Google Scholar and Web of Science were used to search for data on forest elephants feeding preferences using English and French keywords: “forest elephant”, “Loxodonta cyclotis”, “browsing preferences”, “feeding preferences”, “diet”, “diet selection”, “regime alimentaire”, and “elephant de foret”. We also searched recursively through the references provided by the articles that were found during the search. We only retained data from studies that quantified feeding preferences per tree species through ordinal ranking, count of browsing events, selection ratio, or browsing frequency. We excluded studies providing only a list of consumed species. Our data collected in Ndoki were added to the data found in literature for a total of eight studies. Five out of eight studies classified feeding preference/frequency in three categories: rare, medium, high. The Ndoki data contained four categories that were recategorized in three by combining the low and medium-low categories into “low”. The data from Short 1981 reported the number of browsing events per tree species. We assigned species to three categories (low, medium, high browsing preference) based on the frequency distribution of browsing events. Species with less than three browsing events were assigned to the “low” category, species with more than six were assigned to the “high” category, and the species in between to the “medium” category. Feeding preferences in Tchamba&Seme 1993 were reported with an ordinal scale and thus are presented without using categories. Statistical differences between groups were determined using the Student’s t-test for normally distributed data and the Wilcoxon rank-sum test for non-normally distributed data. Dispersal mode of trees was determined following ^6^ and complemented with data collected at LuiKotale ^21^

### Tree inventory data and taxonomy harmonization

Tree inventory data were collected in Ndoki (200 m^2^ plots away from elephant trails, total ∼ 42 ha, 5674 trees DBH > 40 cm) and LuiKotale (16 1-ha plots, 6579 trees DBH >10 cm). In Ndoki, 1664 understory circular plots were enumerated, in which 6479 trees and shrubs were measured and identified. Tree species data (browsing preference plus forest inventories) spanned over several decades and species names were homogenized and updated following the taxonomy provided by World Flora Online and by using their associated R package.

### Wood density data and AGC analysis

We used the R package BIOMASS to assign WD to each feeding record starting from the species level, to the genus and finally to the family. If none of these were available, we assigned the plot-average WD. We group the feeding records in 4 groups following the estimated amount consumed and performed pairwise Student’s two-sided t-tests across the 4 groups to test for statistically significant differences.

Aboveground carbon was calculated using location-specific DBH-height allometries. We simulated the loss of AGC due to the lack of recruitment of Elephant-obligate trees by adapting a previously used methodology^17^. We replaced Obligate trees with new trees which were randomly sampled without replacement from the remaining trees. The relative abundance of the dispersal modes was maintained. This process was repeated 10,000 times for each of the two sites and the difference between pre and post-replacement calculated for each iteration. The mean and standard deviation of the 10,000 iterations were used to estimate the loss of Elephant-obligate trees on AGC.

### Nutritional values of plants

We gathered nutritional values of plant species consumed by elephants from *PNuts*, a global database of plant nutritional values (Berzaghi et al. in prep). *PNuts* contained 102 species of plants part of the forest elephant diet. For each species, data were available for the following nutritional properties: crude protein, nitrogen detergent fiber, acid detergent fiber, dry matter digestibility, condensed tannins, and ash (minerals). These data were complemented with additional data on other leaf and fruit properties that determine food intake quality: sugar, fat, starch, cellulose, and carbohydrates^40^.

### Analysis of effect on vegetation

We have researched the literature using Google Scholar and Web of Science to find studies investigating the physical effect of forest elephant on the ecosystem. The following keywords were used: “forest elephant”, “Loxodonta cyclotis”, “ecosystem engineering”, “ecosystem engineer”, “regeneration”, “mortality”, “tree density”, “stem density”, “debarking”, and “nutrients”. We also examined any relevant publication within the references cited by the articles found during the systematic literature search.

## Acknowledgements

We thank the Governments of the Republic of Congo for collaboration and for permission to conduct elephant ecology research. We are grateful to the African Elephant Conservation Fund of the U.S. Fish and Wildlife Service, the Wildlife Conservation Society, Save the Elephants, United States Agency for International Development (US-AID CARPE), GEFCongo, and the Columbus Zoo Conservation Fund. This study could not have been realized without the astonishing ecological knowledge and forest skills of our tracking team, including Gregoire Mambeleme, Sylvan Imalimo, Mammadou Gassagna, Eric Mossimbo, Zonmimputu and Simon Lamba. Additional technical and logistical assistance was given by G. Kossa Kossa, M. Fay, B. Curran, D. Bourges, Peter D Walsh and Fiona Maisels.

## Author contributions

Conceptualization: FBe, SB, FBr

Methodology: lead by FBe with contributions from SB and CD

Visualization: FBe

Funding acquisition: FBe

Writing: lead by FBe with contributions from all other co-authors

## Data availability

LuiKotale vegetation plot data is available at ForestPlots.net; Ndoki data is available upon request; all other data is available from their respective sources.

