## Supplementary information for "Megaherbivores modify forest structure and increase carbon stocks through multiple pathways"

**This PDF file includes:**

Figs. S1 to S8

Table S1

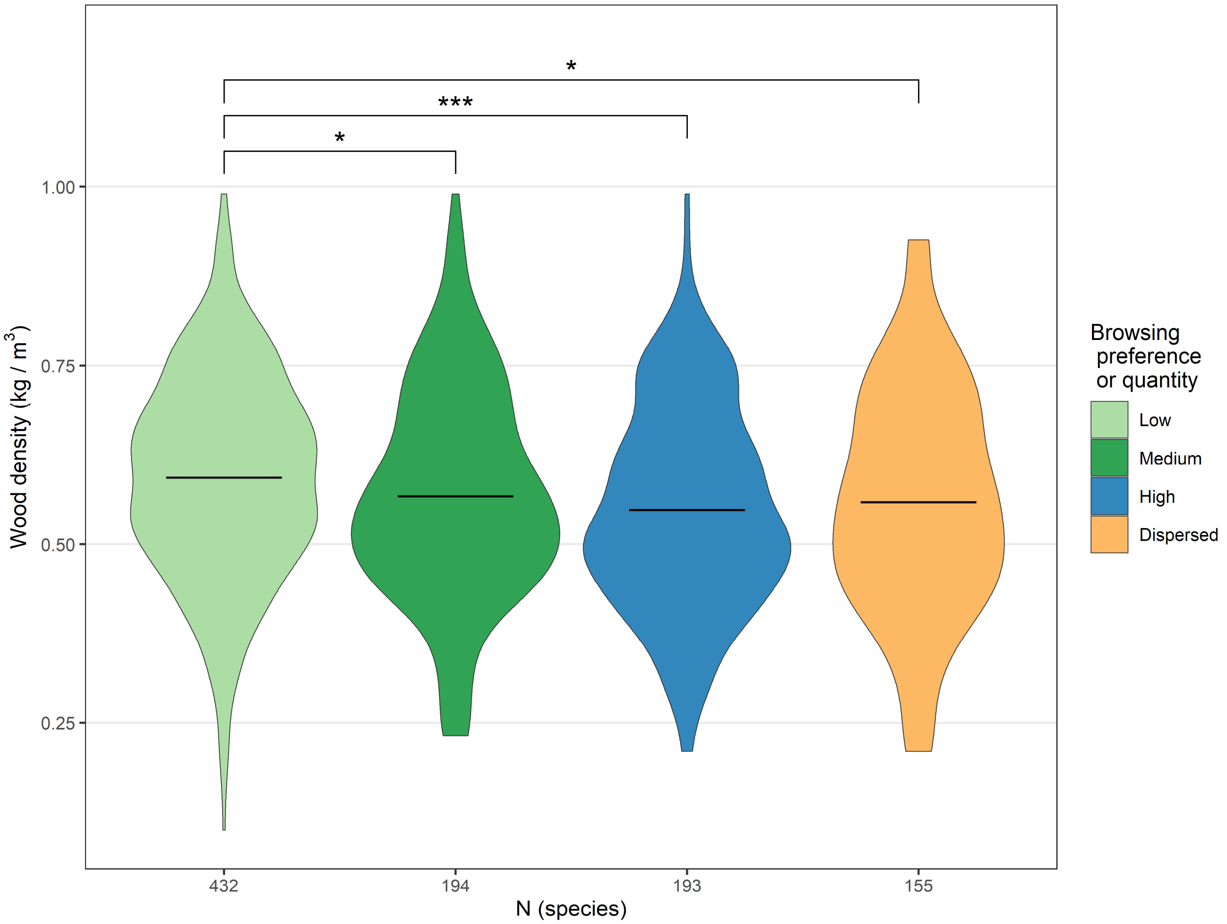


**Fig. S1 Wood density of elephant-dispersed and browsed trees by preference across tropical Africa.** This data does not include browsing preferences from Ndoki. Elephants prefer to browse on low wood density trees and disperse seeds from trees with high wood density. The elephant-dispersed group includes all tree species of which seeds were dispersed by elephants and includes elephant-obligate and non-obligate (dispersed by elephants and other animals). Significance level of pairwise statistical comparison: *P < 0.05; ***P < 0.001.

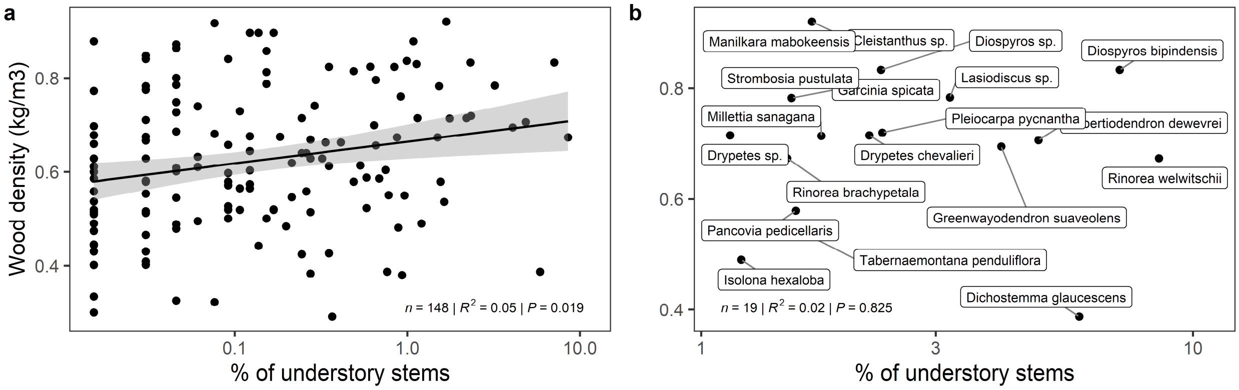


**Fig S2. Species wood density as a function of their abundance in understory plots at Ndoki.** Each point represents a species. (a) All species; (b) selected species constituting 75% of all sampled understory stems.

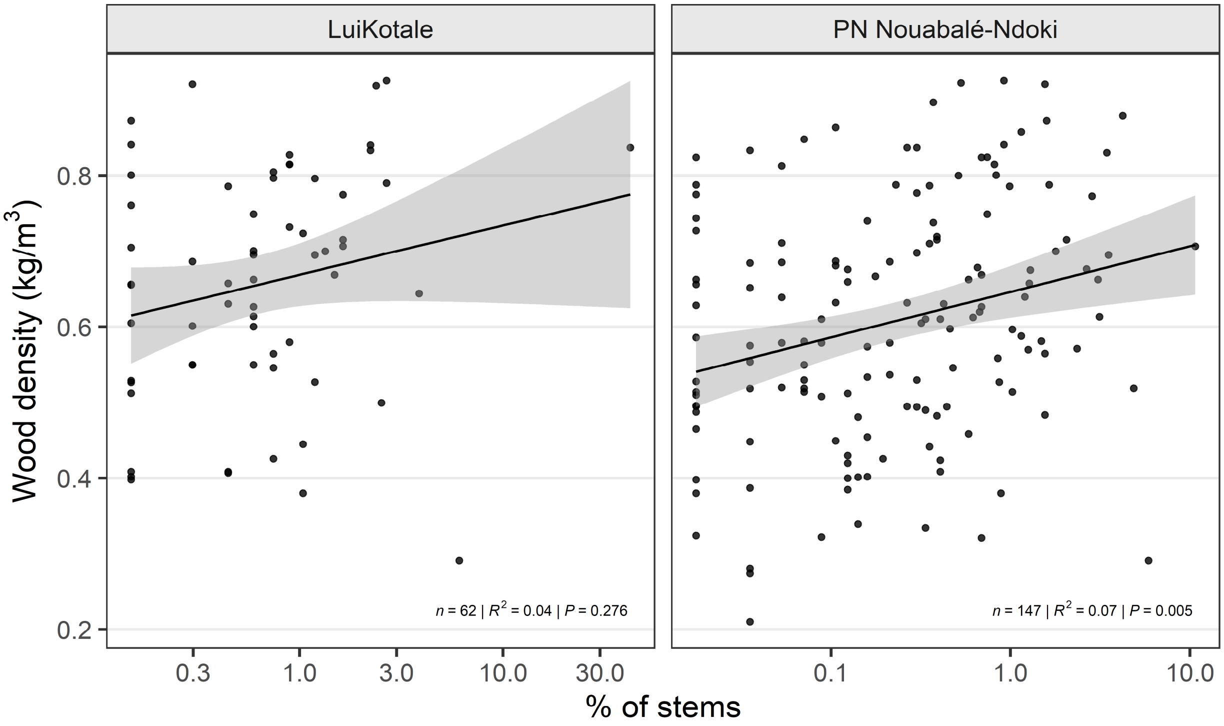


**Fig. S3. Species wood density as a function of their relative abundance in vegetation plots at two sies.** Vegetation plots include all stems with DBH ≥ 40 cm (Methods).

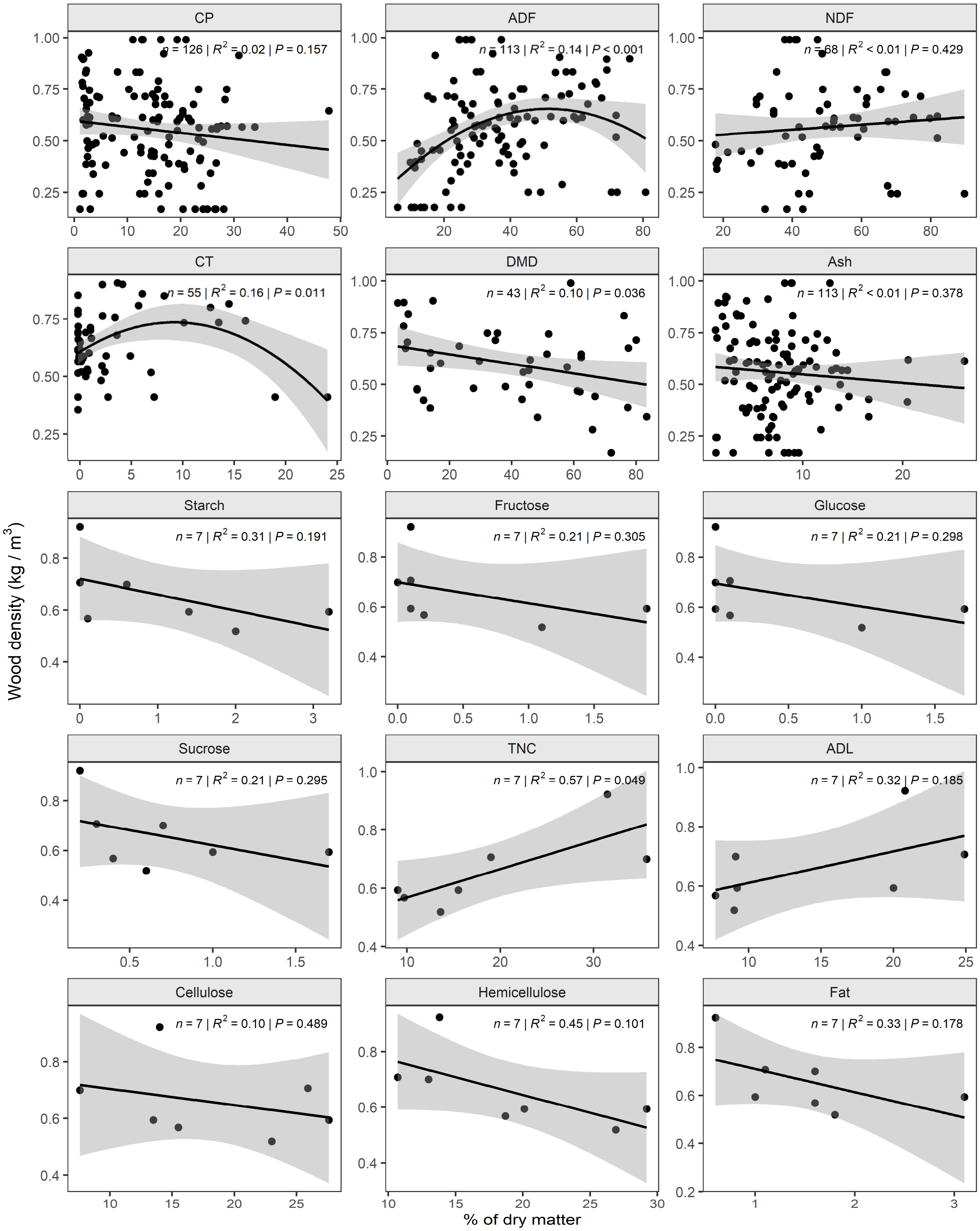


(legend in following page)

**Figure S4.** Correlations between wood density and leaf nutritional properties. CP = crude protein, TNC = total non-structural carbohydrates, DMD = dry matter digestibility, Ash = minerals, NDF = neutral detergent fiber, ADF = acid detergent fiber, ADL = acid detergent lignin, CT = condensed tannins. Data were fitted with a linear model except for ADF and CT which were fitted with a second order polynomial.

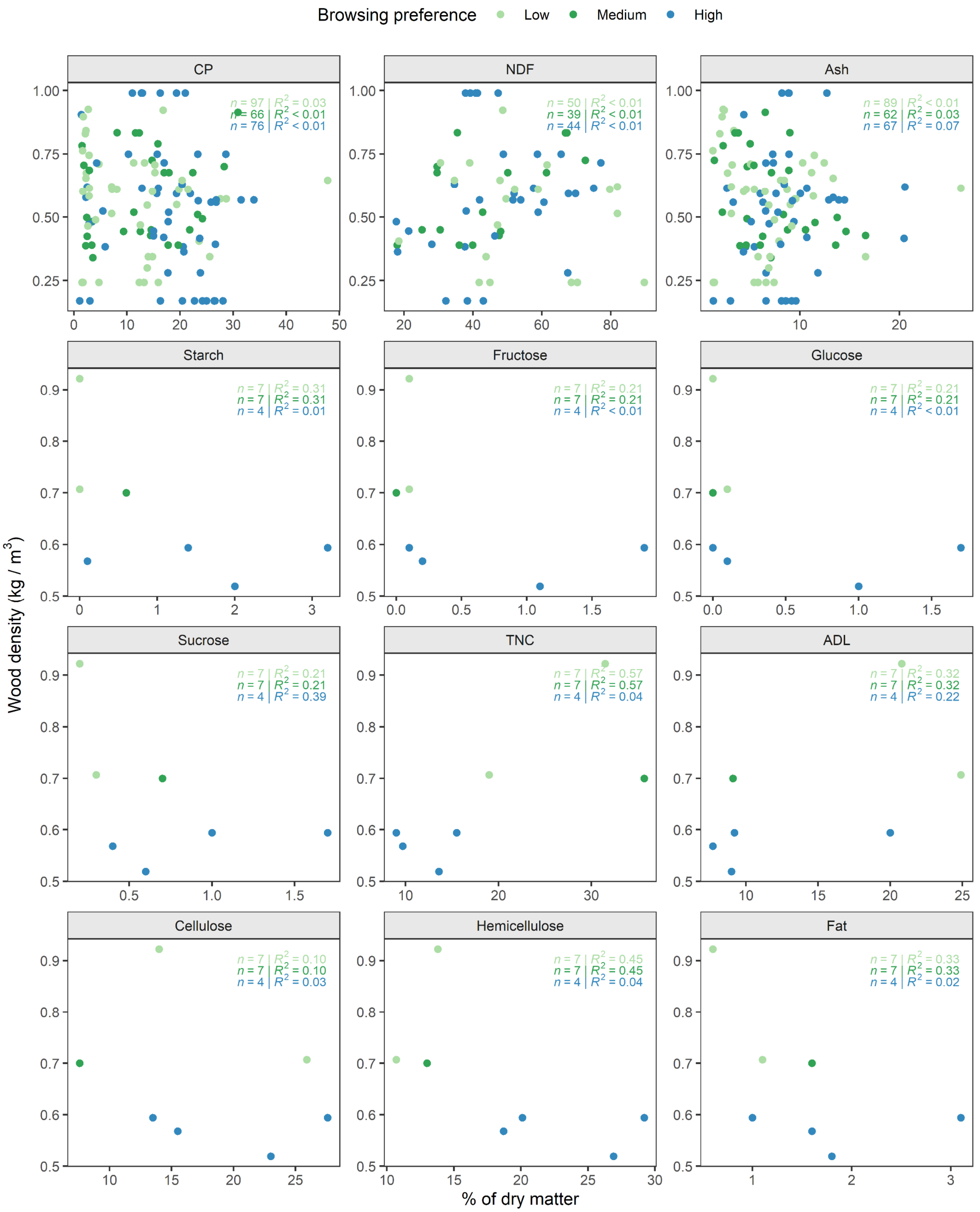


**Fig. S5**. Correlation between wood density and leaf nutritional properties across browsing preference groups. CP = crude protein, TNC = total non-structural carbohydrates, NDF = neutral detergent fiber, ADF = acid detergent fiber, ADL = acid detergent lignin, CT = condensed tannins. For most properties sample size was too small to detect any differences among groups.

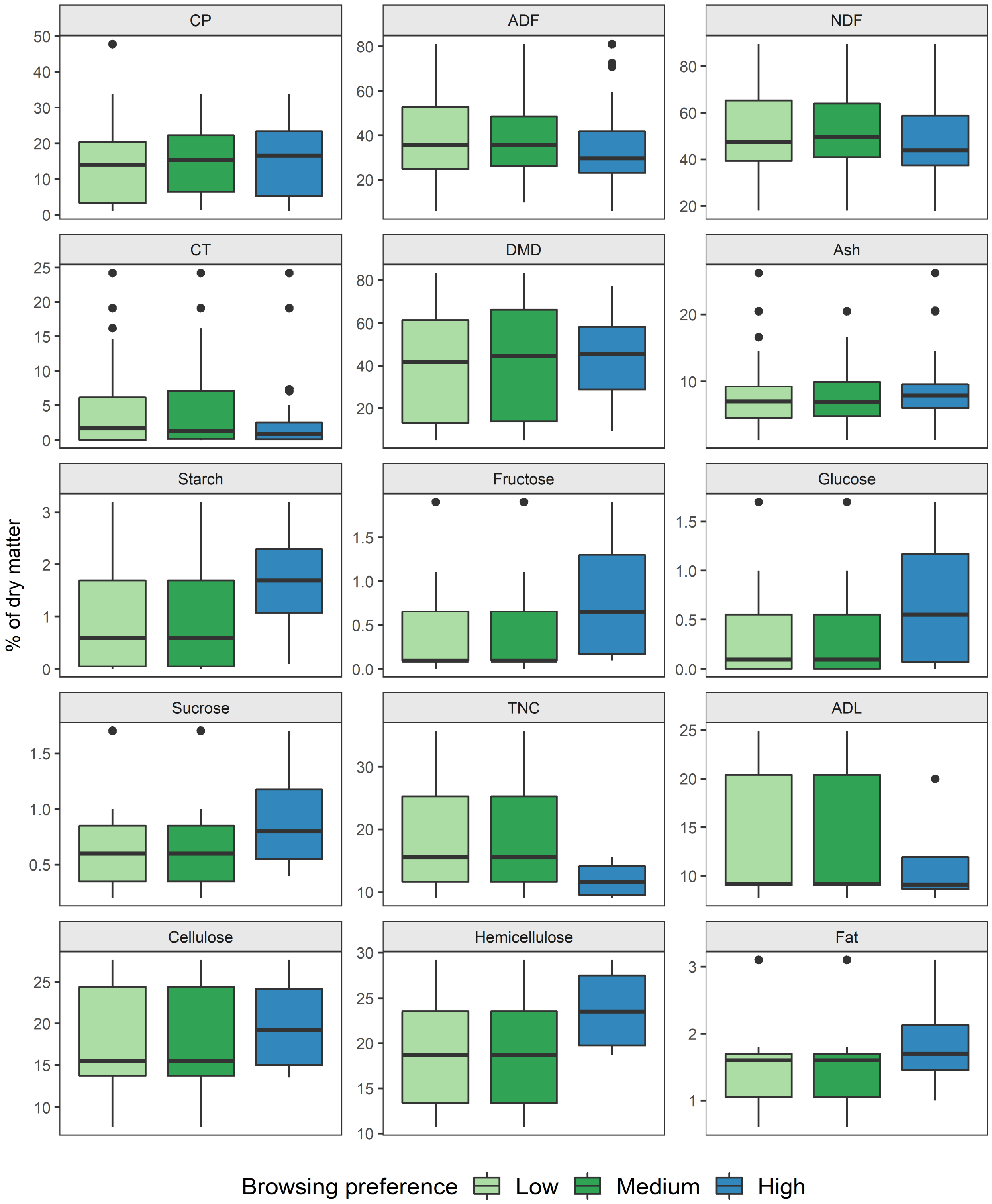


**Fig. S6**. Median, 1st and 3r quartiles of leaf nutritional properties across browsing preference groups. CP = crude protein, TNC = total non-structural carbohydrates, DMD = dry matter digestibility, Ash = minerals, NDF = neutral detergent fiber, ADF = acid detergent fiber, ADL = acid detergent lignin, CT = condensed tannins. No statistically significant differences were detected among the groups (t-test).

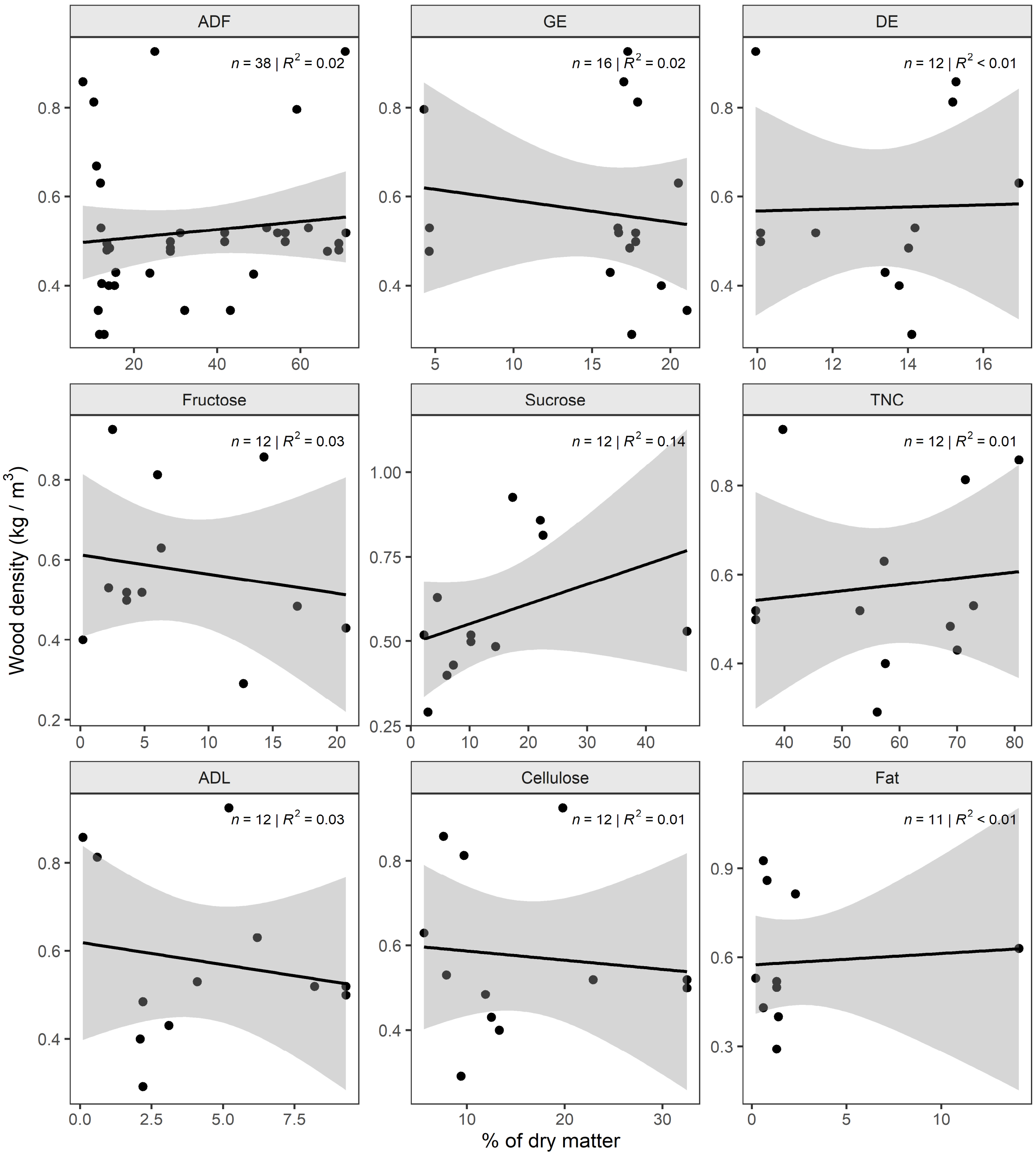


**Fig. S7**. Correlation between wood density and fruit nutritional properties across browsing preference groups. ADF = acid detergent fiber, GE = gross energy, DE = digestible energy, ADL = acid detergent lignin, CT = condensed tannins. Data were fitted with linear a model.

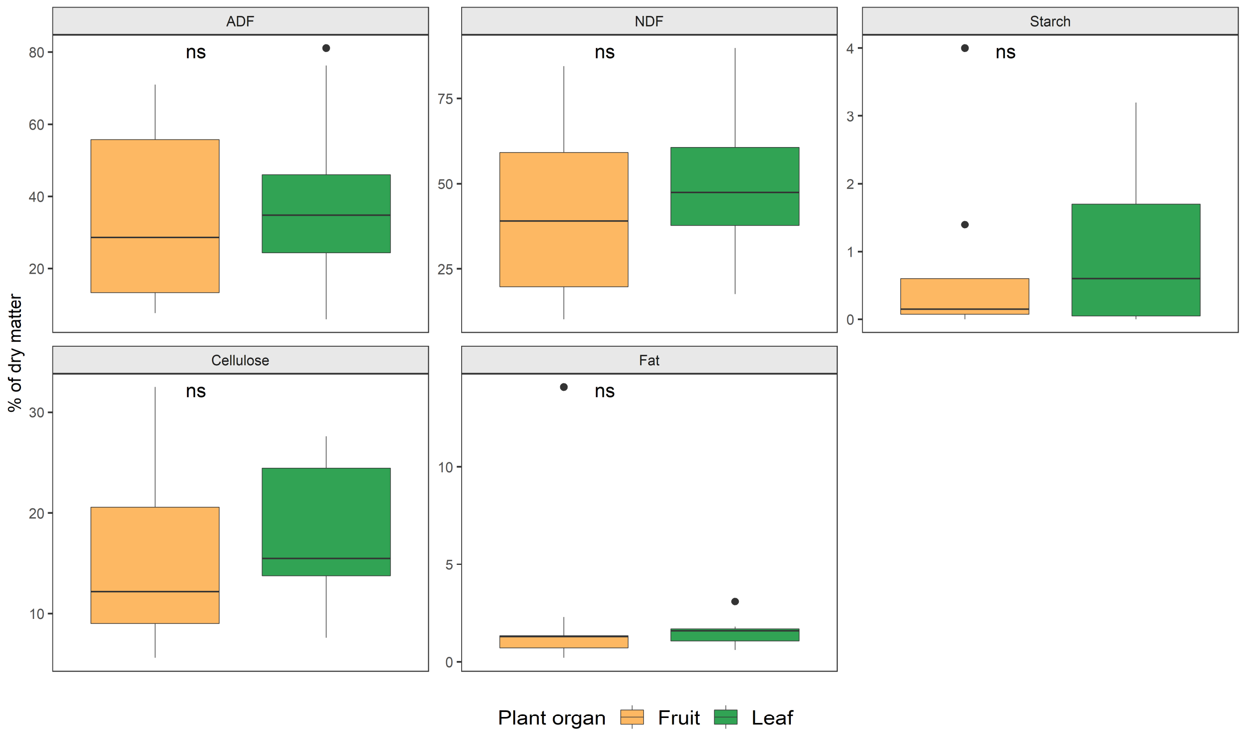


**Fig. S8.** Comparison between fruit and leaf nutritional properties. NDF = neutral detergent fiber andc ADF = acid detergent fiber. Non-statistical difference between the means of the two groups (t-test) are indicated with ‘ns’.

| Author | Site | Country | Lat | Lon | Year | Preference categories | Plants sampled |
|---|---|---|---|---|---|---|---|
| Tchamba&Seme | Santchou | Cameroon | 5.2 | 10 | 1993 | - | n/a |
| Short | Bia | Ghana | 7 | -2.5 | 1981 | (3) | 340 |
| Ouattara | Sassandra | Ivory Coast | 7 | -7 | 2000 | 3 | n/a |
| Merz | Tai | Ivory Coast | 5.6 | -7 | 1981 | 3 | 58,214 |
| Theuerkauf | Bossematie | Ivory Coast | 5.5 | -3.5 | 2000 | 3 | n/a |
| This study | Ndoki | Republic of Congo | 2.7 | 16.5 | 2003 | 4 | 13,211 |
| Wing&Buss | Kibale | Uganda | 0.5 | 30.2 | 1970 | 3 | 118,618 |
| Struhsaker | Kibale | Uganda | 0.5 | 30.2 | 1996 | 3 | 7,174 |

**Table S1. Location of study sites where forest elephant browsing preferences were recorded.** Preference categories in Bia were determined based on the statistical distribution of number of feeding events per species (Methods).
